# Efficient protein structure archiving using ProteStAr

**DOI:** 10.1101/2023.07.20.549913

**Authors:** Sebastian Deorowicz, Adam Gudyś

## Abstract

**Motivation:** The introduction of Deep Minds’ Alpha Fold 2 enabled prediction of protein structures at unprecedented scale. AlphaFold Protein Structure Database and ESM Metagenomic Atlas contain hundreds of millions of structures stored in CIF and/or PDB formats. When compressed with a general-purpose utility like gzip, this translates to tens of terabytes of data which hinders the effective use of predicted structures in large-scale analyses.

**Results:** Here, we present ProteStAr, a compressor dedicated to CIF/PDB as well as, supplementary PAE files. Its main contribution is a novel approach to predict atom coordinates on the basis of the previously analyzed atoms. This allows efficient encoding of the coordinates which are the largest component of the protein structure files. By default, the compression is lossless, though the lossy mode with a controlled maximum error of coordinates reconstruction is also present. Compared to the competing packages, i.e., BinaryCIF, Foldcomp, PDC, our approach offers superior compression ratio at established reconstruction accuracy. By the efficient use of threads at both compression and decompression stages, the algorithm takes advantage of multicore architecture of current central processing units and operates with speeds about 1 GB/s. The presence of C++ API further increases the usability of the presented method.

**Availability and implementation:** The source code of ProteStAr is available at https://github.com/refresh-bio/protestar.

## Introduction

For over 50 years, structural biologists have been collecting atomic-level models of proteins in Protein Data Bank (PDB) [2]. Nowadays it contains more than 200 thousand experimentally-verified protein structures and is an invaluable resource for researchers. Still, this number is orders of magnitude smaller then the number of known proteins. The recent publication of deep learning methods like AlphaFold2 [5] and ESMFold [7] was, therefore, a game changer. These tools can produce high-quality predictions of the 3D structure of a protein in minutes on a workstation. This allows individual researchers to fold the proteins they work on by themselves.

In the large studies, however, involving thousands or even millions of structures, the prediction costs can still be prohibitive. Therefore, the aforementioned utilities have recently been used to construct large databases like AlphaFold Protein Structure Database (APSD) [14] with about 214M predictions or ESM Atlas [7] with 772M entries. This translates to, respectively, 23 TB and 17 TB of gzip archives (this is how the databases are distributed) and a few times larger volumes of uncompressed data. Storing of a local copy of them or even downloading is challenging as well as time and money consuming.

The data in ESM Atlas are provided in textual PDB format [15]. A key component of PDB files and the major contributor to their overall size is a table of Cartesian atom coordinates. The data in APSD are provided in PDB and mmCIF [15] formats. The latter is also textual (with similar atom table) but can contain additional fields. An important feature of AlphaFold2 is that it produces also Predicted Aligned Error (PAE) files storing the accuracy of the relative positions of each pair of atoms. The PAE files are given in JSON format and, when gzipped, contribute similarly as PDBs to the APSD database size.

In the recent years, several attempts have been made to develop a compact representation of PDB and mmCIF files that would outperform general-purpose compressors like gzip. An interesting approach is BinaryCIF [11]. The most important idea behind it is to handle mmCIF tables in a column-wise manner and store numeric fields delta-compressed (i.e., store a difference between the current value and the value in the previous row). These files can be further gzip-compressed to achieve even better compression ratio. In the recent Foldcomp [6] compressor for PDB files, the coordinates of amino acid atoms are stored in records of a fixed size. The algorithm stores bond and torsion angles as well as side chain angles and uses them during the decompression to reproduce atom coordinates. This is space efficient, but the method is inherently lossy. The backbone and sidechain atom positions are reproduced with an average error of ∼80mÅ and ∼140mÅ, respectively, though the maximum error may still exceed 1Å. By redefining the distance between atoms whose positions are stored exactly (25 by default), the user may reduce the reconstruction error at the cost of the compression ratio.

The most recent method, also designed for PDB format, is PDC [16]. In the lossless mode, it stores the coordinates differentially, similarly as BinaryCIF. An interesting idea used here is to reorganize atoms within a residue to minimize the differences between successively encoded entries. In the lossy mode, PDC stores approximate coordinate differences between neighboring C*α* atoms as well as torsion and side chain angles.

Kim *et al*. (2023) evaluated several other approaches to compress protein structures including PULCHRA [8], MMTF [3], PIC [13]. Though, BinaryCIF, Foldcomp, and PDC can be considered as the current state-of-the-art. What is important for huge datasets, only Foldcomp allows storing many files in a single archive and provides fast random access to selected structures. This significantly reduces the maintenance cost as it prevents from keeping and navigating hundreds of millions of separate files at the disk.

In this article, we introduce ProteStAr, a novel compression approach for PDB, mmCIF, and PAE files. It allows storing many files in a single archive and decompress them on demand. For this purpose, we developed two compression components. The first one handles PDB and mmCIF files. A key idea here is to predict coordinates of successive atoms and store only errors of the predictions. As the throughput of the algorithm was of crucial importance, the prediction technique is simple and particularly suited for compression. Thus, it should not be treated as anything comparable to AlphaFold or ESMFold. The second ProteStAr component is PAE compression. Since these files contain square matrices of size equal to the number of atoms in the structure, our algorithm employs some ideas from lossless image compressors.

## Methods

### General overview of the archive format

The ProteStAr archive allows storing many files of the following protein-structure-related formats:

- protein structures in PDB and mmCIF formats,
- predicted align errors in PAE format (produced mainly by AlphaFold),
- confidence files in JSON format containing accuracy of predictions of residues (produced mainly by AlphaFold).

The format can be extended in the future if new file types need to be archived.

The tool is written in the C++17 programming language and is provided as a command-line utility allowing: compression, decompression of all/selected items, listing. Moreover, we provide decompression library for C++ and plan to support also Python in the near future.

Each file is compressed separately, which has some negative consequences in terms of compression ratio, but allows accessing any file in the archive rapidly and easy extension of the existing archives by new entries.

### Compression of PDB and mmCIF files

Since PDB and mmCIF formats are closely related and store similar data, we designed a unified compressor for them differing mainly by a parser component.

Each file is split into sections, which can be a *block* or a *table*. We gather all *block* sections and then compress them using a general-purpose ZSTD compressor. Processing of *tables* is more complicated. First, we determine a table type which could be one of the following: *generic, atom, hetatm*. A table is classified as *atom* if it contains one or more chains of amino acids with typical order of atoms, and without missing fields. Similarly, a table is classified as *hetatm* if it contains information of HETATMs without missing data. The remaining tables are classified as *generic*, which allows handling various types of input files.

A *generic* table is processed in a column-wise manner. For a numeric column (determined on the contents) we check if it is of some special kind, i.e., all values are equal, differences between neighbor rows are the same. If so, the minimum information needed to reconstruct the column is stored. Otherwise, we calculate the differences between the neighbor values (*delta-coding*) and store them using an entropy coder (in particular, a range coder [10]). If the column contains coordinates (but the table is *generic* for any reason) and the user selected lossy compression mode (will be explained in following subsections) we reduce the resolution of the values and encode the differences between neighboring numbers using an entropy coder. The remaining columns are concatenated and ZSTD-compressed.

An *atom* table usually is a major part of PDB/mmCIF file, so efficient compression of it is crucial. Since the order of atoms in every amino acid is established and we know the sequence of amino acids, we do not need to store atom types. The compression of Cartesian coordinates is complex and will be explained in the next subsection. Our tool allows the user to decide if the compression should be lossless (default) or lossy. In the latter case, the user can specify the max. error of position of backbone as well as side chain atoms. If provided, we round the coordinates accordingly (will be described in the next subsection) and handle them losslessly. This is an important advantage over Foldcomp and PDC. They also support lossy compression of coordinates, but the user cannot specify the error bounds and the individual errors could significantly exceed the average.

In the case of AlphaFold2, predictions of the B-factor field are usually constant for a residue. If this is the case, we store these values compactly. Otherwise, we store separate B-factor for every atom, but the user can provide a flag telling the tool that B-factors should be averaged over a residue. B-factors are delta-coded and stored using an entropy coder. The remaining columns in *atom* tables are handled in a similar way like in *generic* tables.

Currently, *hetatm* tables are handled in the same way as *generic* tables, but we plan to implement more sophisticated variants employing some ideas from *atom* sections.

### Compression of atom coordinates

The existing methods, like Foldcomp and PDC, make use of the classic description of how the protein structure is folded. They store angles, which can be used to reconstruct the atom coordinates. For the sake of compactness, the angles are saved with a limited precision making the reconstruction also imprecise. Moreover, the errors can accumulate, which is the main reason, why the algorithms are unable to control the max. error.

Our approach works differently and is free of this disadvantage. We predict the coordinates of each atom making use of the 3 closest, already processed atoms (*reference* atoms). The candidates are backbone atoms from the previous residue as well as the atoms that have been already stored in the current residue (this is why some fixed ordering of atoms in residues is required).

The prediction is done with a use of pre-trained models that were incorporated into the tool. As the training data we used Human proteome from APSD v4 comprising of about 186k mmCIF files. The package contains the learning module, so if necessary, the user can repeat this stage on his/her own mmCIF/PDB files and recompile the tool. Nevertheless, changing the model makes the archives incompatible, so the module is provided mostly for research purposes.

In the first step of training, for each atom in each residue type we calculate the distances to the atoms that could serve as references. Then, for each atom type and each residue type we pick 3 *reference* atom types that are on-average the closest.

In the second step of training, for each atom type of each residue type we collect information about up to *N* = 10^6^ appearances of this atom type. Each observation is stored as a 6-tuple ⟨*r*_12_, *r*_13_, *r*_23_, *f*_1_, *f*_2_, *f*_3_⟩ containing distances between 3 reference residues as well as distances between the *fourth* atom and the references. Each such record can be treated as description of a tetrahedron. We will be using these tetrahedrons to predict atom coordinates at the compression stage.

In the third training step, we perform the clustering of 6-tuples using the well-known k-means algorithm [4]. We try clustering for *k* = 1, …, 20 centroids and pick the value that minimizes the estimated cost of coordinate encoding. The process of selection of *k* mimics (in some sense) what we do in compression, so we will go back to this step after describing how the compression works.

Now, let us assume that we want to encode the coordinates of the *current* atom CE1 in HIS. In the model, we see that the reference atoms are: ND1, CD2, and CG of the current residue. The model contains 3 centroids for this atom type that define 3 tetrahedrons. Now we examine all the tetrahedrons and check which of them allows prediction of the *current* atom position with the lowest error. Conceptually, for each centroid we construct a tetrahedron and then replace its base by changing the distances between the reference atoms to the distances in the current situation, as they are known at both the compression and decompression runs. We keep the distances to the fourth atom as they are in the centroid. Such modified tetrahedrons predict the positions of the *fourth* atom. We compare these predictions with the position of the *current* atom and select the best one. An illustration of this process is given in Figure 1.

**Fig. 1.**
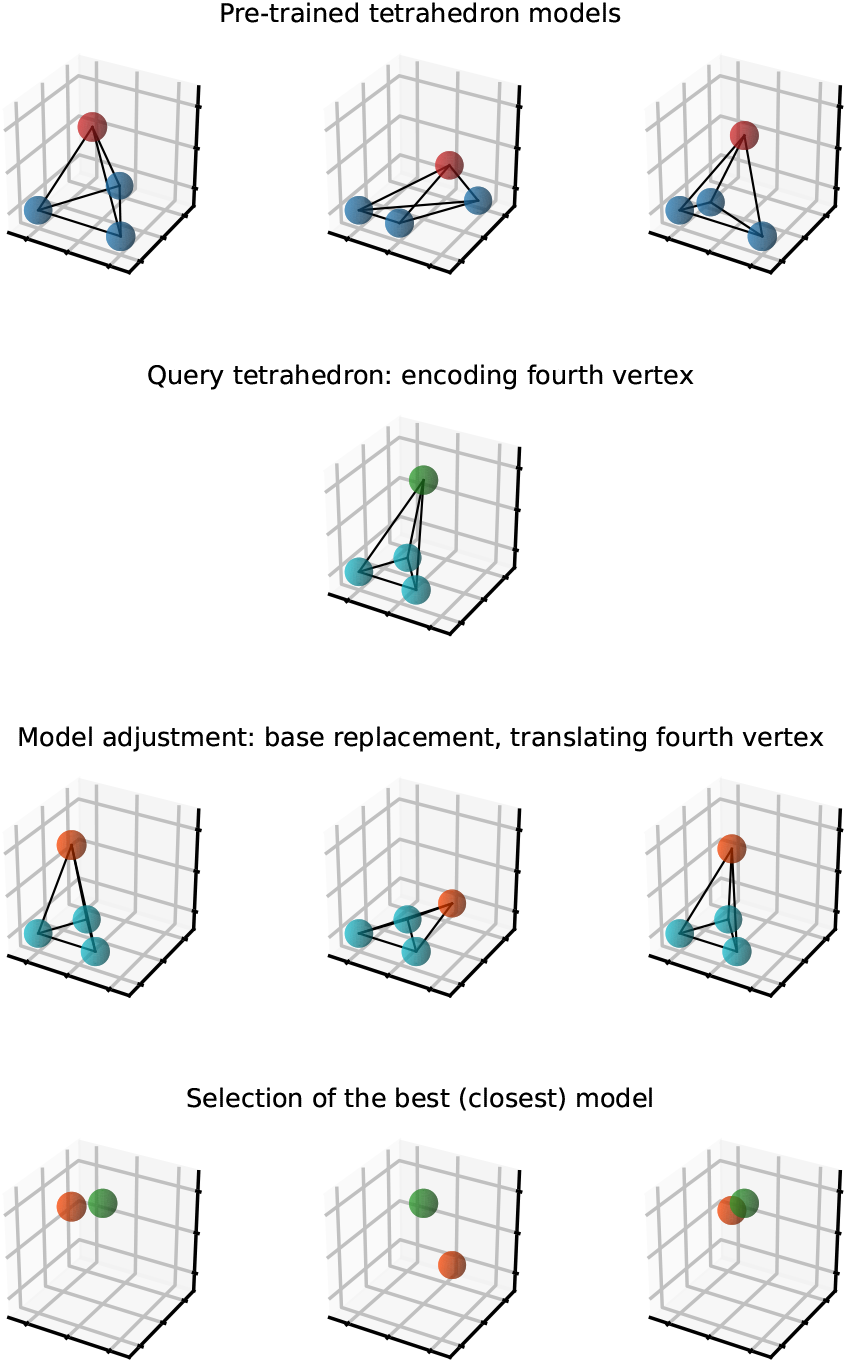
Illustration of how the prediction of the current atom works. First, we take from the model the centroids describing tetrahedrons. Second, we calculate a tetrahedron for the current situation. Third, we use the distances of the current atom to the reference atoms to adjust tetrahedrons from the model. Fourth, we calculate the distances between the predictions of the position of the current atom given by the model and the current one to find the best prediction.

Technical details are a bit more complicated. First, the tetrahedrons are not *normalized* (i.e., the first reference atom is not at (0, 0, 0)). We use the real coordinates of the reference atoms in the current situation and construct tetrahedrons in the real space. Then, we estimate the encoding cost of the prediction error as:

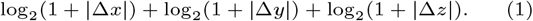

Thus, it may happen that we will select worse prediction (i.e., the larger Euclidean distance) if it could be encoded cheaper.

After selection of the best centroid, we entropy-encode its identifier, Δ*x*, Δ*y*, Δ*z*, and the identifier of the tetrahedron (for each centroid we can construct two equivalent tetrahedrons and we check both). If replacing the base makes it impossible to construct a tetrahedron with preserved distances to the fourth atom, the positions of the first two reference atoms are returned as predicted positions. Handling of the first three atoms of a chain is special: as there are no reference atoms for them, their coordinates are delta-encoded.

In the lossy mode, the user provides the max. error for backbone atoms and, separately, for side chain atoms. Then, prior to processing we round the input coordinates of all atoms following the formula:

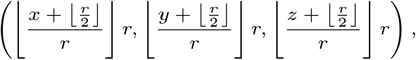

Where 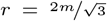 and *m* is the max. error. Then, we prune the model for each atom type by removing the centroids that are identical after coordinates rounding, which can potentially reduce the cost of encoding centroid id.

We provide also a simplified selection of a centroid for prediction. Instead of examining all of them we just calculate the 6-tuple for the current atom and look for the closest centroid in the model (Euclidean distance is used here). This is significantly faster but leads to a bit worse compression ratio. The user can select this optional mode during the compression.

Now, let us go back to the learning stage. When we decide which value of *k* (the number of centroids) to use, we try to estimate the encoding cost of the *N* collected observations using the current model. The cost is estimated using the formula (1) increased by the cost of encoding the centroid id: log_2_(1 + *k*). We pick the value of *k* that minimizes the estimated cost of encoding. In our model, the value of *k* varies between 1 (N in HIS) and 20 (CG in GLU).

### Compression of PAE files

Predicted align errors produced by AlphaFold2 tell how trustworthy are the relative positions of every pair of atoms. Since a file stores a square matrix containing integer values from the range [0, 32], our approach employs some ideas from the lossless image compression methods like PNG [9], FLIF [12], JPEG-XL [1].

We process the matrix in a row-wise manner and column-wise within a row. We predict the value of the current item *v*(*i, j*) taking into account the neighboring items. Let us denote the prediction as *e*(*i, j*). First, we decide if the current item should be predicted horizontally or vertically. To do this, we check statistics (built on the already processed part of the matrix) how frequently the difference between an item and its upper neighbor was smaller in the *j*-th column than between an item and its left neighbor in the *i*-th row. We choose the direction that more likely leads to the smaller difference. Without loss of generality, let us assume that *e*(*i, j*) is predicted horizontally (the other case is symmetric). If *i* = *j*, we predict 0, since this is PAE value for the same atom. If |*i* − *j*| = 1, we predict 1. Otherwise we check if *v*(*i, j* − 1) − *v*(*i* − 1, *j*) *>* 4. If so we predict *e*(*i, j*) = *v*(*i, j* − 1) − 1. If not and *v*(*i* − 1, *j*) − *v*(*i, j* − 1) *>* 4 we predict *e*(*i, j*) = *v*(*i, j* − 1) + 1. If not we use *e*(*i, j*) = *v*(*i, j* − 1).

We encode the error of prediction using an entropy coder in some context which also depends on the neighborhood. The context is combined from the following:

- *v*(*i* − 1, *j*) − *v*(*i* − 1, *j* − 1) quantized to 5 values (less than −1, −1, 0, 1, more than 1),
- *v*(*i, j* − 1) − *v*(*i, j* − 2) quantized to 5 values in the same manner,
- *v*(*i, j* − 2) − *v*(*i, j* − 3) quantized to 3 values (less than 0, 0, more than 0),
- *v*(*i, j* − 1) − *v*(*i* − 1, *j*) quantized to 3 values,
- *v*(*i* − 1, *j* − 1) − *v*(*i* − 1, *j* − 2) quantized to 3 values,
- *v*(*i, j* − 1) quantized to 6 values ([0, 7], [8, 26], [27, 28], 29, 30, [31, 32]),
- information that we predicted horizontally.

Then, we entropy-encode the error of prediction, i.e., *e*(*i, j*) − *v*(*i, j*) in this context. There are 8100 possible contexts which allows storing the statistics in the fast cache memory making the algorithm fast. Slightly better compression ratios would be possible for large matrices at the cost of more complicated model and slower processing.

As for some purposes, the full resolution of PAE files is not necessary, we propose also a lossy scheme. The processing is the same with the only important difference being ‘rounding’ the original values according to the scheme presented in Table 1 and small differences in how the context is determined.

**Table 1.** Grouping of PAE values for supported lossy levels. Every value from [*ℓ, r*] range is reconstructed during decompression as 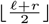.

| Level | Grouping |
| --- | --- |
| 1 | 0, 1, ..., 19, 20–21, 22–23, 24–27, 28–32 |
| 2 | 0, 1, ..., 9, 10–12, 13–15, 16–19, 20–23, 24–27, 28–32 |
| 3 | 0, 1, ..., 5, 6–7, 8–10, 11–14, 15–19, 20–25, 26–32 |
| 4 | 0, 1, 2, 3, 4–5, 6–8, 9–12, 13–17, 18–24, 25–32 |

### Compression of other file types

At the current version of the tool we also allow to store confidence files in JSON format. Such files are small as they contain only 3 arrays of size equal to the number of residues. They contain: residue id (integer), confidence score (float value), confidence category (single character). Thus, we just ZSTD-compress them.

The archive format is, however, open, so it is possible add other file types in the future.

### Experimental results

In the experiments, we used two large databases of predicted protein structures: APSD v4 and ESM Atlas containing 214M and 772M structures, respectively. Testing machine was equipped with 64-core AMD 3995WX Pro CPU clocked at 2.7 GHz and 512 GiB RAM running under openSUSE Tumbleweed operating system. The disks were: NVME Seagate FireCuda 530 4 TB and RAID5 composed of four Seagate Exos 16 TB HDDs. If not stated otherwise, the experiments were carried out on the NVME disk. Details on the datasets, tools, command-lines used in this study are given in the Supplementary Material.

As a first step, we evaluated the performance of the tools on 542k AlphaFold predictions of SwissProt proteins as well as a subset of 59k predictions of various qualities from ESM Atlas database. Foldcomp and ProteStAr, as the only multi-threaded packages in the analysis, were run with 16 computing threads. Since gzip and BinaryCIF are one-to-one compressors, the collection of their output files (one per structure) was gathered using well-known tar utility. This was to avoid space overhead related to storing on disk hundreds of thousands of files. The summary of the experiments is presented in Figure 2. Detailed results are given in the Supplementary Worksheet.

**Fig. 2.**
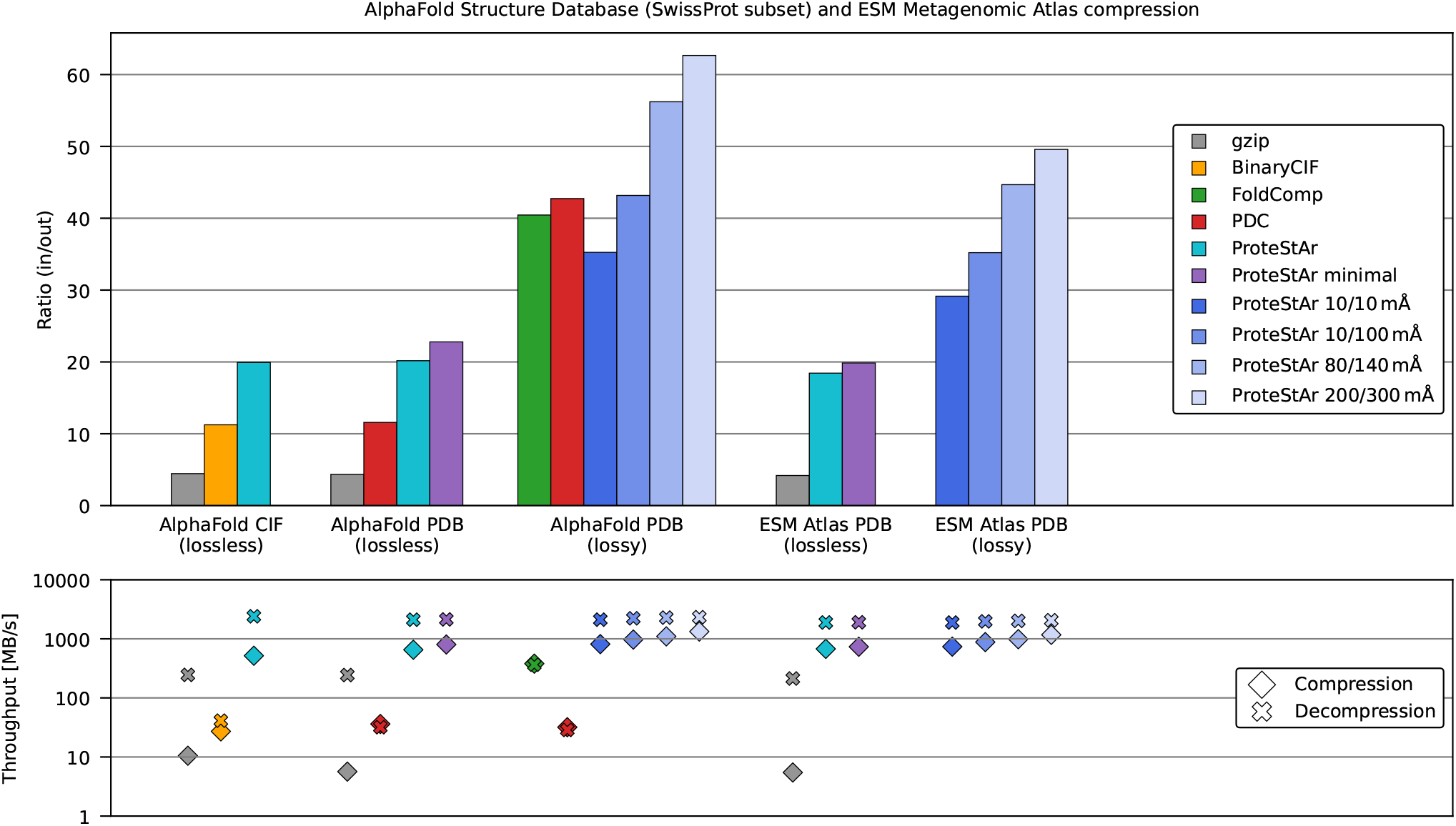
Comparison of CIF/PDB compressors for subsets of APSD and ESM Atlas databases. Compression ratios are calculated as input_size/compressed_size. Series `ProteStAr` *x*/*y* represent lossy variant of our algorithm with max. error of *x* mÅ for backbone atoms and *y* mÅ for side chain atoms.

When considering lossless compression, our algorithm was 4 times better then gzip and 2 times better then state-of-the-art BinaryCIF (for mmCIF files) and PDC (for PDB files). Storing only essential parts of PDB files (the header and the table with atom descriptions) in ProteStAr minimal mode further increased the compression ratio. The compression and decompression speeds of our tool were about 500 MB/s and 2400 MB/s, respectively—significantly higher then those of the competitors.

For the lossy compression, ProteStAr was compared to PDC and Foldcomp. As these tools store only header and atom table of PDB files, our tool was configured in the same way. Moreover, ProteStAr, unlike competitors, allows controlling max. error of atom coordinates reconstruction. Therefore, we investigated several lossy variants of our algorithm with error rates comparable to the competing tools. In particular, series `ProteStAr 80/140` with the error bounds equal to 80 mÅ and 140 mÅ for backbone and side chain atoms, respectively, was selected as a direct competitor of Foldcomp. Analogously, `ProteStAr 200/300` was compared with PDC. As the results on AlphaFold SwissProt show, both these variants outperformed state-of-the-art methods in terms of compression ratio. What is interesting, even `ProteStAr 10/100` mode reduced the size of the input data by a factor of 43, which is roughly ten times better than gzip. This shows how much space can be saved when using a specialized tool and allowing reduction of precision of atom coordinates. Unfortunately, we were not able to evaluate any of the competitors on 59k subset from ESM Atlas. PDC failed to process all the files, while Foldcomp produced invalid archives for files inconsistent with the PDB format specification (e.g., missing atoms, errors in labeling of residues), which are present in ESM Atlas. The results of ProteStAr were, however, similar to those observed for AlphaFold SwissProt predictions.

In the next experiment, we evaluated the error of atom coordinates reconstruction of Foldcomp, PDC, and selected lossy modes of ProteStAr. The analysis relied on compressing and decompressing all AlphaFold predictions of mouse proteome (21,615 files) and comparing the reconstructed atom positions with the input ones.

The resulting error histograms of backbone and side chain atoms, together with compression ratios are presented in Figure 3a. As one can observe, the reconstruction accuracy of ProteStAr was always within the requested bounds, while the errors of PDC and Foldcomp were sometimes significantly larger than the average.

**Fig. 3.**
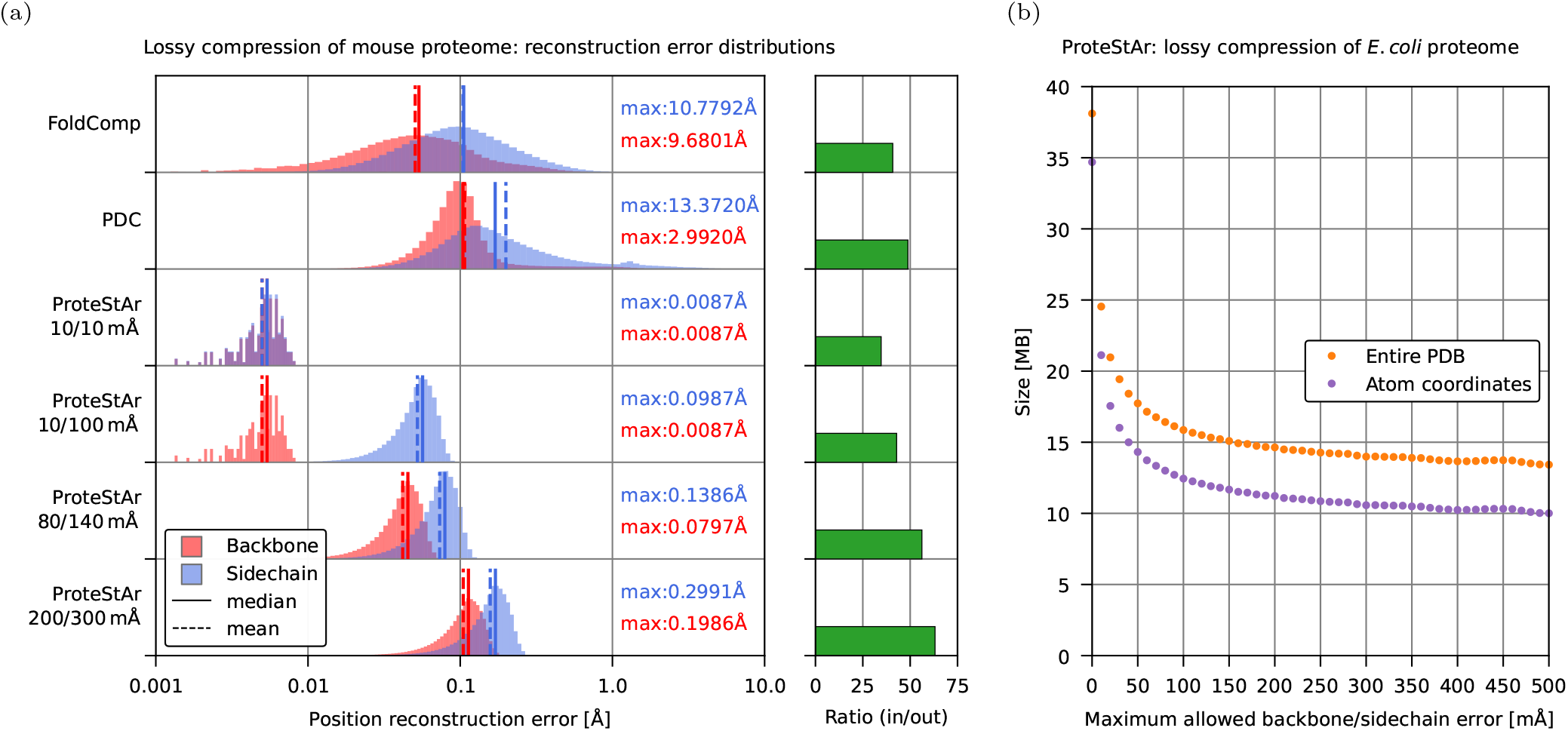
Evaluation of error of atom coordinates. (a) Histograms of errors for backbone (red) and side chain (blue) atom positions. Median, average, and maximum errors are also presented. Green bars show compression ratio (input_size/compressed_size). (b) Impact of maximum error of atom positions (same for backbone and side chain) on the overall archive size (orange) and size of the coordinates alone (purple).

As a next step we measured the impact of max. allowed reconstruction error on ProteStAr compression ratio. The results for APSD *E*.*coli* dataset, containing 10.5 million atoms (from 2500 randomly selected files), are presented in Figure 3b. As one can see, the file size rapidly dropped from 38.1 MB for lossless compression to 15.9 MB for max. errors of 100 mÅ and then the improvement became moderate. This can be, partially, explained by the fact that when the coordinates are rounded, the accuracy of the prediction of the fourth atom position decreased. Nevertheless, for 500 mÅ error bound the archive size was 13.4 MB with approximately 10 MB used for atom coordinates. This translates to less then 8 bits for storing 3D coordinates of a single atom.

Table 2 shows the sizes of the archives produced by the examined tools. We used 5 proteomes from AlphaFold database, SwissProt proteins, some subset of ESM Atlas and the full ESM Atlas (version v0 and the supplement). The largest dataset was used to show the scalability of the examined tools. ProteStAr, depending on the mode, needed from 29 to 31 hours to complete for ESM Atlas v0. What is also important, the ProteStAr compression ratios were almost independent on the dataset, in spite of training internal models on human AlphaFold predictions. All the investigated packages are memory efficient—the maximal observed memory usage in the experiments were 7 MB for gzip, 12 MB for PDC, 170 MB for Foldcomp, and 311 MB for ProteStAr.

**Table 2.** Comparison of compressed sizes of the tools for 5 proteomes of model organisms, SwissProt proteins, and ESM Atlas. The largest dataset is split into: initial (v0) release and the supplement (v2023_02).

| Dataset | No. struct. | Raw [MB] | gzip [MB] | PDC lossless [MB] | PDC Lossy [MB] | Foldcomp lossy [MB] | ProteStAr lossless [MB] | ProteStAr minimal [MB] | ProteStAr 10/10 [MB] | ProteStAr 80/140 [MB] |
| --- | --- | --- | --- | --- | --- | --- | --- | --- | --- | --- |
| E.coli | 4,363 | 878 | 203 | 77 | 22 | 22 | 44 | 38 | 25 | 16 |
| Human | 23,391 | 9,579 | 2,176 | 792 | 184 | 234 | 459 | 427 | 277 | 170 |
| Maize | 39,299 | 9,865 | 2,276 | 844 | 222 | 245 | 490 | 439 | 286 | 178 |
| Mouse | 21,615 | 7,068 | 1,615 | 592 | 145 | 173 | 343 | 314 | 204 | 126 |
| Yeast | 6,039 | 1,913 | 436 | 161 | 40 | 47 | 93 | 85 | 55 | 34 |
| Swissprot | 542,378 | 124,628 | 28,677 | 10,766 | 2,916 | 3,081 | 6,179 | 5,469 | 3,535 | 2,217 |
| ESM Atlas subset | 58,539 | 7,006 | 1,674 | — | — | — | 380 | 353 | 240 | 157 |
| ESM Atlas v0 | 592,006,392 | 67,704,502 | 15,498,908 | — | — | — | 3,777,870 | 3,503,725 | 2,419,293 | 1,592,709 |
| ESM Atlas v2023.02 | 49,059,840 | 7,558,311 | 1,689,967 | — | — | — | 415,442 | 392,043 | 270,068 | 178,533 |

In the final experiment, we evaluated the compression of PAE files. We are not aware of any specialized compressor for this format, so we compared ProteStAr with gzip, which was employed for storing PAE in AlphaFold database. Since PAE files are useless if not accompanied with PDB/mmCIF structures, we used the entire APSD human proteome composed of 186,016 triples of files: (i) mmCIF describing a structure, (ii) PAE showing the accuracy of relative position of every pair of atoms, (iii) confidence of the prediction of each residue. The results are presented in Figure 4. As one can observe, in the lossless mode, ProteStAr compressed PAE files 2 times better than gzip, while in the lossy schemes the advantage over gzip was much larger. When we focus on the size of the whole archive, we can see that the initial dataset of size 119.2 GB could be packed in the lossless mode about 3 times better than gzip, to just 7.5 GB. This drops to 2.5 GB if we decide to the most aggressive scheme in which atom coordinates are stored with 200/300 mÅ max. error and the PAE value resolution is reduced from 33 to 11 distinct values. The compression and decompression speeds of PAE files ranged from 800 MB/s (lossless mode) to 950 MB/s (lossy modes). When we consider all file types together, these values were 600–790 MB/s in compression and 1090–1270 MB/s in decompression. The memory footprint of ProteStAr was about 2 GB in compression and 1 GB in decompression.

**Fig. 4.**
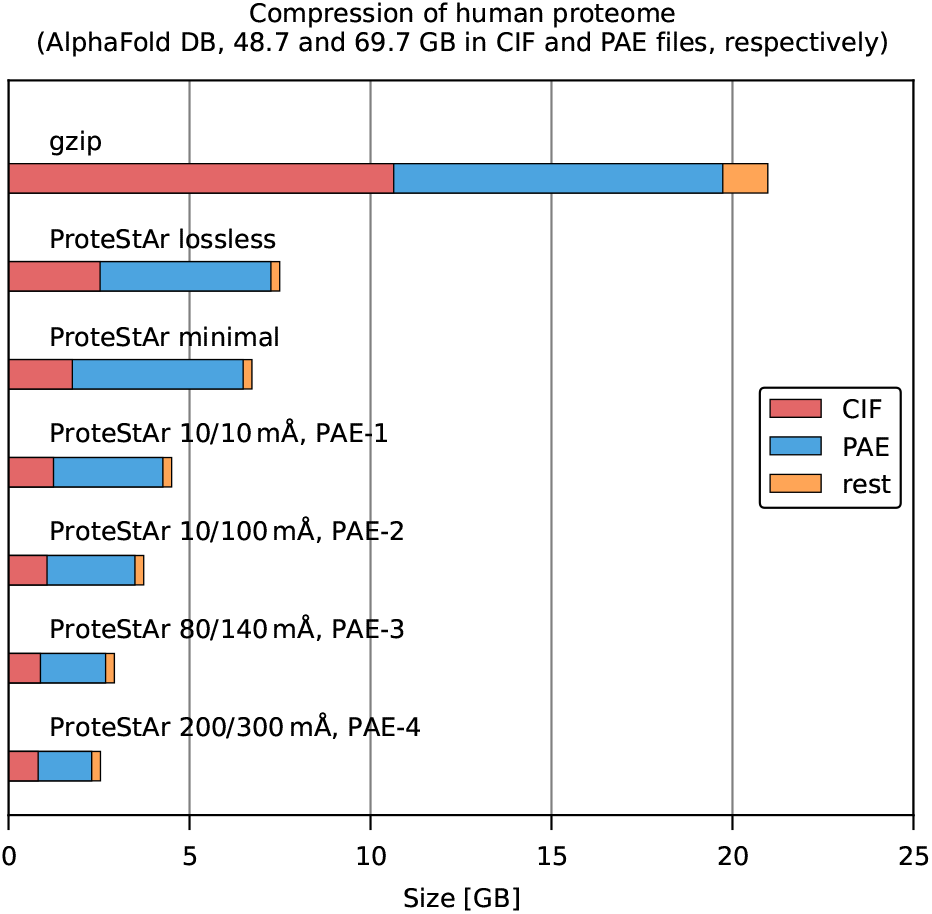
Compression of the entire human proteome (186,018 proteins, 119.2 GB in total). There are 3 files for each protein: mmCIF (48.7 GB in total), PAE (69.7 GB), confidence (0.7 GB). Various lossy schemes are presented for both, CIF and PAE files.

The same APSD human proteome dataset (558,054 files in total, 7.5 GB in ProteStAr lossless or 21.0 GB in tar) was used to check random access speed of ProteStAr. The results highly depended on where the archive resided (NVME, HDD), how large was the file compared to the RAM of the machine and whether the file was cached by OS (due to previous queries) or not. To evaluate this we performed the experiments in three scenarios: (i) file at NVME and cached by OS, (ii) file at NVME, not cached, (iii) file at HDD, not cached. For ProteStAr the time to list the archive contents was less than a second in all examined cases. For tarred archive the times were, respectively, 2.8 s, 2.9 s, 14.4 s. Extraction of a single file by ProteStAr took from 0.02 s to 0.25 s depending on the file size and these times were almost the same in the examined scenarios. The times for tar were: 1.0 s, 13.2 s, 31.6 s, respectively. Clearly, using tar was significantly more time consuming, especially when data were not cached. Alternatively, one can store gzips as they are (without tar) allowing fast random access to the data, at the cost of dealing with hundreds of millions of files, which can be problematic. As Foldcomp does not support PAE and confidence files, we were not able to include it in this comparison. Nevertheless, for databases with PDB/mmCIF structures, the extraction of a single file was similarly fast as for ProteStAr.

## Conclusions

With a rapidly increasing size of databases of protein structure predictions, the general-purpose gzip would soon become insufficient in terms of both, the compression ratio and the processing speed. Our tool, ProteStAr, is able to compress mmCIF and PDB files losslessly four times better then gzip and two times better then state-of-the art BinaryCIF and PDC algorithms. If this reduction is not enough, one can use lossy compression with a controlled error of position reconstruction. This feature, not provided by any of the existing algorithms, allows compacting the data approximately ten times better then gzip, while maintaining maximum reconstruction error at very good levels: 10 mÅ for backbone and 100 mÅ for side chain atoms. All this is obtained at compression/decompression rates comparable to disk throughput (∼1 GB per second) with very fast random access to the archive entries. The functionality of the software is complemented by the support of non-structure files distributed in the prediction databases (PAE, confidence).

While ProteStAr is resistant to the inconsistencies with the mmCIF/PDB format specification—unlike the competitors, it was able to analyze the entire ESM Atlas database, the current release lacks support of several standard elements (e.g., ANISOU and SIGUIJ entries in PDB files are skipped). This issues will be fixed in the future versions.

## Supporting information

Supplementary Worksheet

## Author contributions statement

S.D. designed and implemented the majority of the algorithm. A.G. contributed to the algorithm and implementation. S.D. designed and performed the experiments. S.D. wrote the majority of the manuscript. A.G. contributed to the manuscript. A.G. designed and prepared the visualizations. All authors read and approved the final manuscript.

## References

1. Alakuijala J, et al. JPEG XL next-generation image compression architecture and coding tools. Proceedings Volume 11137, Applications of Digital Image Processing XLII; 111370K (2019).

2. Berman HM, Westbrook J, Feng Z et al. The Protein Data Bank. Nucleic Acids Res 2000 28(1):235–42.

3. Bradley AR, Rose AS, Pavelka A et al. MMTF—An efficient file format for the transmission, visualization, and analysis of macromolecular structures. PLoS Comput Biol 2017;13:e1005575.

4. Forgy E. Cluster Analysis of Multivariate Data: Efficiency versus Interpretability of Classifications. Biometrics 1965;21:768–780.

5. Jumper J, Evans R, Pritzel A et al. Highly accurate protein structure prediction with AlphaFold. Nature 2021;596:583–589.

6. Kim H, Mirdita M, Steinegger M. Foldcomp: a library and format for compressing and indexing large protein structure sets. Bioinformatics 2023;39(4):btad153.

7. Lin Z, Akin H, Rao R et al. Evolutionary-scale prediction of atomic-level protein structure with a language model. Science 2023;379(6637):1123–1130.

8. Rotkiewicz P, Skolnick J. Fast procedure for reconstruction of full atom protein models from reduced representations. J Comput Chem 2008;29(9):1460–1465.

9. Salomon D, Motta G. Handbook of data compression Springer, 5th edition (2010).

10. Schindler M. A fast renormalization for arithmetic coding. a poster in the Data Compression Conference, 1998 available at http://www.compressconsult.com/rangecoder.

11. Sehnal D, Bittrich S, Velankar S et al. BinaryCIF and CIFTools-Lightweight, efficient and extensible macromolecular data management. PLoS Comput Biol 2020;16:e1008247.

12. Sneyers J, Wuille P. FLIF: Free lossless image format based on MANIAC compression. Proceedings of the IEEE International Conference on Image Processing, 2016:pp. 66–70.

13. Staniscia L, Yu YW. Image-centric compression of protein structures improves space savings. bioRxiv 2022, DOI:10.1101/2022.01.20.477098.

14. Varadi M, Anyango S, Deshpande M et al. AlphaFold protein structure database: massively expanding the structural coverage of protein-sequence space with high-accuracy models. Nucleic Acids Res 2021;50:D439–D444.

15. Westbrook JD, Young JY, Shao C et al. PDBx/mmCIF Ecosystem: Foundational Semantic Tools for Structural Biology. JMol Biol 2022;434:167599.

16. Zhang C, Pyle AM. PDC: a highly compact file format to store protein 3D coordinates Database, 2023;2023:baad018

