## Supplementary Material for "Efficient protein structure archiving using ProteStAr"

Sebastian Deorowicz Adam Gudyś

July 20, 2023

**3 Environment** **7**

**4 Additional results** **8**

### 1 Datasets

We used datasets from AlphaFold Protein Structures Database v.4 and ESM Atlas v0 and v2023_02.

#### 1.1 Model organisms proteomes from APSD

Data downloaded from `https://alphafold.ebi.ac.uk/download`.

- Budding yeast
  `https://ftp.ebi.ac.uk/pub/databases/alphafold/latest/UP000002311_559292_YEAST_v4.tar`
- *E.coli*
  `https://ftp.ebi.ac.uk/pub/databases/alphafold/latest/UP000000625_83333_ECOLI_v4.tar`
- Human
  `https://ftp.ebi.ac.uk/pub/databases/alphafold/latest/UP000005640_9606_HUMAN_v4.tar`
- Maize
  `https://ftp.ebi.ac.uk/pub/databases/alphafold/latest/UP000007305_4577_MAIZE_v4.tar`
- Mouse
  `https://ftp.ebi.ac.uk/pub/databases/alphafold/latest/UP000000589_10090_MOUSE_v4.tar`
- Swiss-Prot CIF
  `https://ftp.ebi.ac.uk/pub/databases/alphafold/latest/swissprot_cif_v4.tar`
- Swiss-Prot PDB
  `https://ftp.ebi.ac.uk/pub/databases/alphafold/latest/swissprot_pdb_v4.tar`

#### 1.2 Human proteome (large)

The variant of human proteome with PAE and confidence files was downloaded from Google Cloud Public Datasets as described at: `https://alphafold.ebi.ac.uk/download`.

#### 1.3 ESM Atlas — full datasets

The full datasets are downloaded from `https://github.com/facebookresearch/esm/tree/main/scripts/atlas`. We followed the links from `https://github.com/facebookresearch/esm/blob/main/scripts/atlas/v0/full/tarballs.txt` (v0 version) and `https://github.com/facebookresearch/esm/blob/main/scripts/atlas/v2023_02/full/tarballs.txt` (v2023_02 version).

#### 1.4 ESM Atlas — subset

For evaluation purposes we picked approx. 59k PDB files from ESM Atlas. Details how to download this dataset are given at: `https://github.com/refresh-bio/protestar`.

### 2 Examined programs

The following programs were used in the experimental part. Running parameters are also given.

#### 2.1 gzip v. 1.12

- Compression of a directory

```
# <dataset_name> <in_dir> <out_dir> <tmp_dir>
for i in $2/*; do mv "$i" $4; done
gzip -k -9 $4/*.cif
tar -cf $3/$1.tar $4/*.gz
```

- Decompression of an archive

```
# <dataset_name> <in_dir> <out_dir> <tmp_dir>
tar -xf $3/$1.tar
gzip -d $4/*.gz
```

#### 2.2 BinaryCIF, CIFTools v. 5.0.0

- Compression — we implemented a short script for bulk compression of many CIF files:

```
package org.rcsb.cif;

import org.rcsb.cif.model.CifFile;
import org.rcsb.cif.model.FloatColumn;
import org.rcsb.cif.schema.StandardSchemata;
import org.rcsb.cif.schema.mm.AtomSite;
import org.rcsb.cif.schema.mm.MmCifBlock;
import org.rcsb.cif.schema.mm.MmCifFile;

import java.io.IOException;
import java.net.URL;
import java.util.Optional;
import java.util.OptionalDouble;
import java.util.OptionalInt;
import java.io.File;
import java.nio.file.Path;
import java.nio.file.Paths;

public class CompressBatch {
    public static void main(String[] args) throws Exception {
        // Get args
        // args[0] = input directory that contains cif files
        // args[1] = output directory that will contain bcif files
        Path input = Paths.get(args[0]);
        Path output = Paths.get(args[1]);
        // Get all files in input directory
        File[] files = input.toFile().listFiles();
        // Loop through all files

        for (File file : files) {
            // Get file name
            String fileName = file.getName();
            // Get file extension
            String fileExtension = fileName.substring(fileName.lastIndexOf(".") + 1, fileName.length());
            String outputFileName = fileName.substring(0, fileName.lastIndexOf(".")) + ".bcif";
            // Check if file is a cif file
            if (fileExtension.equals("cif")) {
                // Get input file path
                Path inputFilePath = Paths.get(input.toString(), fileName);
                // Get output file path
```

```
                Path outputFilePath = Paths.get(output.toString(), outputFileName);
                // Measure running time
                long endTime;
                long startTime = System.nanoTime();
                // Handle exceptions
                try {
                    // Read cif file
                    CifFile cifFile = CifIO.readFromPath(inputFilePath);
                    // Write bcif file
                    CifIO.writeBinary(cifFile, outputFilePath);
                    endTime = System.nanoTime();
                    System.out.println(fileName + "\t" + (endTime - startTime) / 1000000000.0);
                } catch (Exception e) {
                    System.out.println(fileName+ "\t" + "NA");
                }
            }
        }
    }
}
```

- Decompression — we implemented the following script

```
package org.rcsb.cif;

import org.rcsb.cif.model.CifFile;
import org.rcsb.cif.model.FloatColumn;
import org.rcsb.cif.schema.StandardSchemata;
import org.rcsb.cif.schema.mm.AtomSite;
import org.rcsb.cif.schema.mm.MmCifBlock;
import org.rcsb.cif.schema.mm.MmCifFile;

import java.io.IOException;
import java.net.URL;
import java.util.Optional;
import java.util.OptionalDouble;
import java.util.OptionalInt;
import java.io.File;
import java.nio.file.Path;
import java.nio.file.Paths;

public class DecompressBatch {
    public static void main(String[] args) throws Exception {
        // Get args
        // args[0] = input directory that contains bcif files
        // args[1] = output directory that will contain cif files
        Path input = Paths.get(args[0]);
        Path output = Paths.get(args[1]);
        // Get all files in input directory
        File[] files = input.toFile().listFiles();
        // Loop through all files
        for (File file : files) {
            // Get file name
            String fileName = file.getName();
            // Get file extension
            String fileExtension = fileName.substring(fileName.lastIndexOf(".") + 1, fileName.length());
            String outputFileName = fileName.substring(0, fileName.lastIndexOf(".")) + ".cif";
            // Check if file is a cif file
            if (fileExtension.equals("bcif")) {
                // Get input file path
                Path inputFilePath = Paths.get(input.toString(), fileName);
                // Get output file path
                Path outputFilePath = Paths.get(output.toString(), outputFileName);
                // Measure running time
                long endTime;
                long startTime = System.nanoTime();
                // Handle exceptions
                try {
                    // Read bcif file
```

```
                    CifFile cifFile = CifIO.readFromPath(inputFilePath);
                    // Write cif file
                    CifIO.writeText(cifFile, outputFilePath);
                    endTime = System.nanoTime();
                    System.out.println(fileName + "\t" + (endTime - startTime) / 1000000000.0);
                } catch (Exception e) {
                    System.out.println(fileName+ "\t" + "NA");
                }
            }
        }
    }
}
```

##### 2.3 Foldcomp v. 0.0.5

- Compression of a directory

```
# <dataset_name> <in_dir> <out_dir> <tmp_dir>
./utils/foldcomp compress -t 16 $2 --db -y $3/$1.fcz
```

- Decompression of an archive

```
# <dataset_name> <in_dir> <out_dir> <tmp_dir>
./utils/foldcomp decompress -t 16 $3/$1.fcz $4
```

##### 2.4 PDC commit c49393d (Mar 7, 2023)

- Compression of a directory—lossless

```
# <dataset_name> <in_dir> <out_dir> <tmp_dir>
for f in $2/*.cif; do
./utils/pdc $f $4/$(basename $f).pdc -l=0
done
tar -cf $3/$1.pdc_lossless_tar $4/*.pdc
```

- Compression of a directory—lossy

```
# <dataset_name> <in_dir> <out_dir> <tmp_dir>
for f in $2/*.cif; do
./utils/pdc $f $4/$(basename $f).pdc -l=2
done
tar -cf $3/$1.pdc_lossy_tar $4/*.pdc
```

- Decompression of an archive—lossless

```
# <dataset_name> <in_dir> <out_dir> <tmp_dir>
tar -xf $3/$1.pdc_lossless_tar
for f in $4/*.pdc; do
./utils/pdd $f $4/$(basename $f).cif -f=2
done
```

- Decompression of an archive—lossy

```
# <dataset_name> <in_dir> <out_dir> <tmp_dir>
tar -xf $3/$1.pdc_lossy_tar
for f in $4/*.pdc; do
./utils/pdd $f $4/$(basename $f).cif -f=2
done
```

##### 2.5 ProteStAr v. 0.7.0

- Compression of a directory—lossless

```
# <dataset_name> <in_dir> <out_dir> <tmp_dir>
./utils/psarch add --type pdb -t 16 --indir $2 --out $3/$1.psarch_pdb_lossless -v 0
```

- Compression of an archive—minimal

```
# <dataset_name> <in_dir> <out_dir> <tmp_dir>
./utils/psarch add --type pdb -t 16 --indir $2 --out $3/$1.psarch_minimal -v 0 --minimal
```

- Compression of an archive—10/10

```
# <dataset_name> <in_dir> <out_dir> <tmp_dir>
./utils/psarch add --type pdb -t 16 --indir $2 --out $3/$1.psarch_minimal_10_10 -v 0
    --minimal --lossy --max-error-bb 10 --max-error-sc 10
```

- Compression of an archive—10/100

```
# <dataset_name> <in_dir> <out_dir> <tmp_dir>
./utils/psarch add --type pdb -t 16 --indir $2 --out $3/$1.psarch_minimal_10_100 -v 0
    --minimal --lossy --max-error-bb 10 --max-error-sc 100
```

- Compression of an archive—80/140

```
# <dataset_name> <in_dir> <out_dir> <tmp_dir>
./utils/psarch add --type pdb -t 16 --indir $2 --out $3/$1.psarch_minimal_80_140 -v 0
    --minimal --lossy --max-error-bb 80 --max-error-sc 140
```

- Compression of an archive—200/300

```
# <dataset_name> <in_dir> <out_dir> <tmp_dir>
./utils/psarch add --type pdb -t 16 --indir $2 --out $3/$1.psarch_minimal_200_300 -v 0
    --minimal --lossy --max-error-bb 200 --max-error-sc 300
```

- Decompression of an archive

```
# <dataset_name> <in_dir> <out_dir> <tmp_dir>
./utils/psarch get --type ALL --all -t 16 --in $3/$1.psarch_pdb_lossless --outdir $4/ -v 0
```

##### 2.6 tar v. 1.34

### 3 Environment

The machine used in the tests was of the following configuration:

- AMD 3995WX Pro CPU clocked at 2.7 GHz CPU, 64 cores,
- 512 GiB RAM,
- NVME Seagate FireCuda 530 4 TB—used in majority of tests,
- RAID5 composed of four Seagate Exos 16 TB HDDs—used in compression of full ESM Atlas,
- openSUSE Tumbleweed operating system.

Our tool was compiled using gcc 11.3.0.

#### 4 Additional results

Additional results are given in the Supplementary Worksheet.
